# Possible Self-awareness in Wild Adélie Penguins *Pygoscelis adeliae* ^¶^

**DOI:** 10.1101/2022.11.04.515260

**Authors:** Prabir Ghosh Dastidar, Azizuddin Khan, Anindya Sinha

**Affiliations:** Polar Science Division, Ministry of Earth Sciences, Government of India, New Delhi, India; SGT University, Gurugram, Delhi NCR, India; Psychophysiology Laboratory, Humanities and Social Sciences, Indian Institute of Technology Bombay, Mumbai, India; Animal Behaviour and Cognition Programme, National Institute of Advanced Studies, Bengaluru, India; Centre for Neuroscience, Indian Institute of Science, Bengaluru, India; Department of Environmental Sciences and Wildlife Biology, Cotton University, Guwahati, India

**Author notes:** **Corresponding authors:** Anindya Sinha, Prabir Ghosh Dastidar. We dedicate this paper to the fond memory of the late David Walton of the British Antarctic Survey, United Kingdom, who showed keen interest in this work but did not wait to see it in its final form.

**Keywords:** Group-behaviour test, modified mirror test, hidden-head test, coloured-bib test, Svenner Island, East Antarctica

## Abstract

This preliminary study, conducted in January–February 2020, investigates the potential presence of self-awareness in a population of wild Adélie penguins on the Dog’s Neck Ice Shelf and on Svenner Island in East Antarctica. It is based on the responses and reactions of individual penguins to images, generated in mirrors during three experimental paradigms: a group-behaviour test; a modified mirror test and a hidden-head test. We believe that this set of experiments constitutes possibly the first investigations into the potential presence of self-awareness in any penguin species and is pioneering in conducting a set of cognitive experiments on free-ranging individuals of a nonhuman species in its natural environment, without any prior familiarisation, conditioning or acclimatisation to the experimental paradigms employed. Future studies, integrating the socioecology and cognitive ethology of penguins, may provide insights into whether our experimental paradigms could provide evidence to confirm the presence of self-awareness and even of self-recognition in this species and examine whether the observed social awareness may have evolved due to the social needs of individual penguins to engage in cooperative behaviour with conspecific individuals, while maintaining their independent decision-making capacities, throughout their communal lives.

## 1. Introduction

Penguins are ancient flightless birds, mostly available in the Southern Hemisphere, and one of the very few species surviving on earth for over 60+ million years, even having outlived the dinosaurs. Amongst the 17 species of penguins found worldwide, only seven species inhabit Antarctica and the sub-Antarctic islands, and of these, the Adélie penguin *Pygoscelis adeliae* is the most abundant, though found exclusively only in Antarctica (Croxall and Prince 1979). They are social birds, with remarkable adaptive features, which include feeding, swimming, nesting and breeding colonially in discrete homogeneous groups, but only in ice-sea zones. The extreme climate of Antarctica necessitates a high level of cooperative behaviour amongst individual penguins to withstand climatic hardship, even leading to the development and maintenance of rookeries, where individuals huddle together for thermoregulatory purposes and exhibit colonial breeding at favourable times of the year.

The test of mirror self-recognition is a behavioural technique, devised by Gallup Jr. (1970) to examine whether particular nonhuman individuals—chimpanzees, in his case— possessed the capacity of self-awareness, as monitored by their ability to recognise themselves as individuals in a mirror (reviewed in Reiss and Morrison 2017). In the original experiments, four chimpanzees, after two days of isolation prior to the test, were individually exposed for eight hours on each of two days to their reflected image. Following another eight days— approximately 80 hours—of prolonged exposure to their reflected images in mirrors, all four chimpanzees, marked with a red dye on a part of their body that they could not directly see, attempted to touch the marks on their own bodies after seeing their reflections in the mirror. This was interpreted as clear evidence that the chimpanzees were self-aware, as they seemed to identify the individuals, bearing the red marks, in the mirrors as being themselves or, in other words, they were capable of self-recognition, as evidenced by this particular paradigm.

Over the years, mirror self-recognition has been reported as an emergent phenomenon, in the absence of explicit training, in several nonhuman species, including, amongst others, the great apes, bottlenose dolphins, Asian elephants and most recently, a fish – the cleaner wrasse (Kohda *et al*. 2022). The comprehensiveness of this test has, however, often been challenged, especially with respect to animals that rely on sensory modalities other than vision to perceive their environment and themselves (Bekoff and Sherman 2004). More generally, should a failure to recognise oneself or pass the mirror self-recognition test necessarily mean that an animal lacks self-awareness or the capacity to become the object of one’s own attention (Safina 2015; Cazzolla Gatti *et al*. 2021)?

In birds, self-awareness has been investigated successfully, using the mirror test, in pigeons (Epstein *et al*. 1981), magpies (Prior *et al*. 2008) and Indian house crows (Buniyaadi *et al*. 2020). Each of the three pigeons, tested by Epstein and his colleagues, for example, used a mirror to locate a spot on its body, which it could not see directly, following independent exposure and habituation to a mirror for less than 15 hours over a 10-day training period. Each of the individual birds had, however, had a variety of prior laboratory experience, similar to Gallup Jr.’s chimpanzees, and they all acquired repertoires similar to those of the experimental chimpanzees. It must be noted, however, that the ability of birds to generally recognise themselves visually in a mirror remains equivocal with jackdaws (Soler *et al*. 2014) and carrion crows (Brecht *et al*. 2020; Vanhooland *et al*. 2020) having failed the mirror mark test (see Brecht and Nieder 2020 for a review).

In this preliminary study, possibly the first one of its kind in any penguin species, we examine the responses and reactions of wild Adélie penguins *Pygoscelis adeliae*, either individually or as a group, to their self-images, generated in a mirror during several field experimental paradigms, including a group-behaviour test, a modified mirror test and a hidden-head test. Our investigations lead us to tentatively suggest that Adélie penguins are possibly self-aware, as indicated by their responses to their own images in a mirror and argue that such self-awareness may play a critical role during the various communal activities and social interactions that individuals of this avian species typically engage in.

## 2. Materials and Methods

### 2.1 Study species and study sites

Our explorations of penguin self-awareness was conducted on either groups of or on individual Adélie penguins *Pygoscelis adeliae*, the most abundant of all penguin species in Antarctica (Croxall and Prince 1979). The various, well-organised, group behaviours that involved coordinated movement, displayed by the species in the extreme climate conditions of Antarctica, made them ideal subjects for natural experiments that required significant sample sizes of experimental subjects.

All natural observations and experiments were conducted on the Dog’s Neck Ice Shelf (S 70°03’43”, E 24°51’58”) and on Svenner Island (S 69°08’12”, E 76°44’45”), both in East Antarctica, during the 39^th^ Indian Expedition to Antarctica in the months of January and February 2020. Svenner Island has a rookery, at a height of about 30 m, bordering the sea, which is home to around 3500 Adélie penguins, making it an ideal site for our experiments.

### 2.2 Familiarisation with the study individuals

In the initial, exploratory phase of our study on the Dog’s Neck Ice Shelf, two of us (PGD and AK), dressed in polar gear, knelt low to observe the natural responses of a group of 17 Adélie penguins, wandering about 800 m away on a vast icesheet, to the crouching humans. Possibly being attracted by the novelty of the situation, the penguin group approached us on their own. The penguins milled around us for a predetermined duration of 15 min, during which time we videographed them for about 2:38 min. The penguins appeared to be comfortable and showed no signs of stress in our presence (Figure 1). This experience gave us the confidence that we could conduct our experiments in the natural habitat of the penguins, without subjecting the test individuals to any prior familiarisation, conditioning or acclimatisation to the experimental paradigms that we would employ.

**Figure 1.**
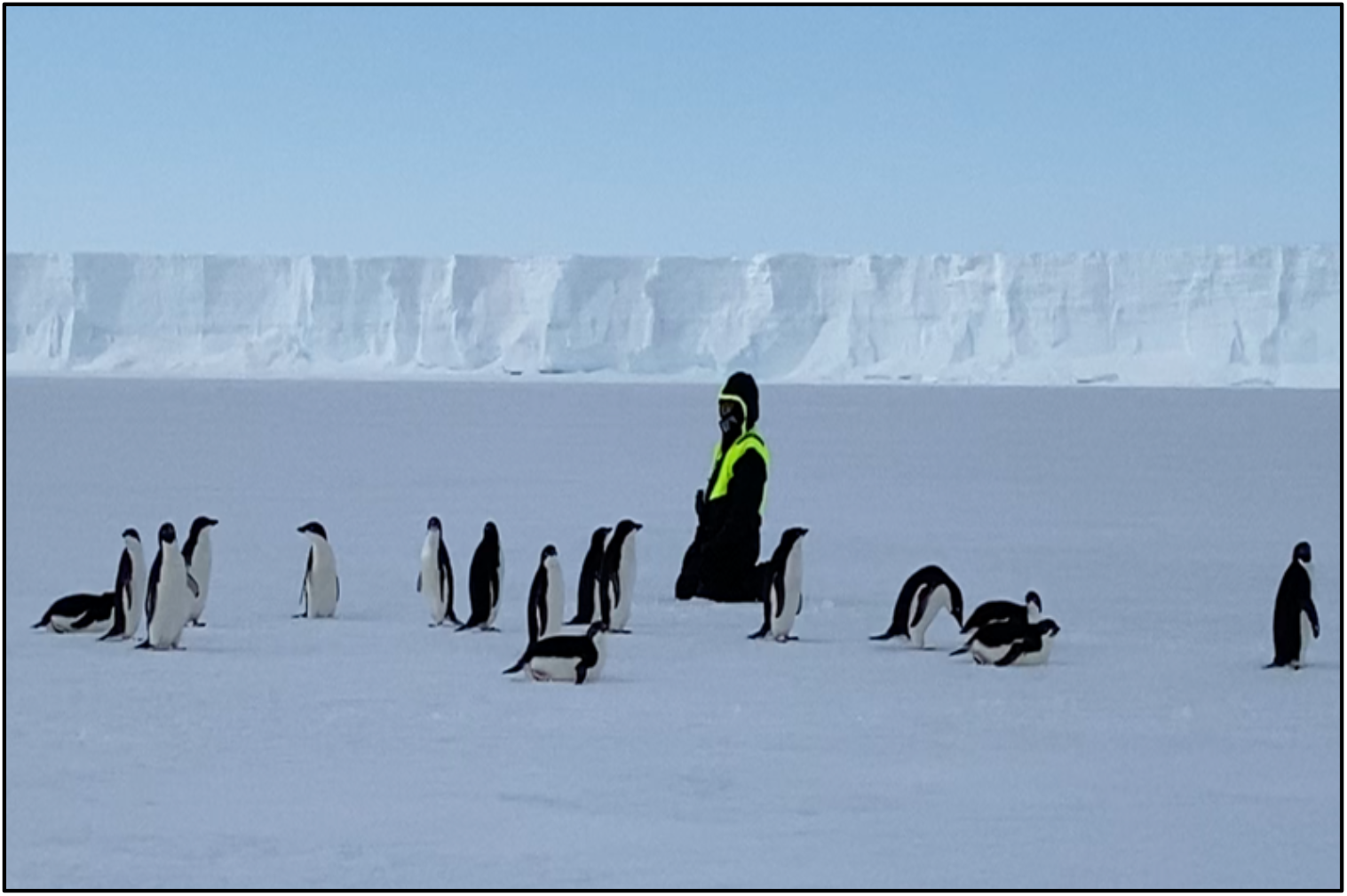
Familiarisation with the study Adélie penguins on Dog’s Neck Ice Shelf, East Antarctica

### 2.3 Experimental paradigms

#### 2.3.1 The group-behaviour test

On Svenner Island, we set up a mirror of approximately 355 × 235 cm in dimension in the path of waddles of Adélie penguins, to observe their reactions, both individually and as a group, to the mirror images that were generated (Figure 2). The mirror that we provided was large enough to be visible from a long distance and attracted relatively large numbers of penguins. On each of the five trials conducted, the mirror was kept in position indefinitely, until it did not attract any penguin for a prolonged period of time.

**Figure 2.**
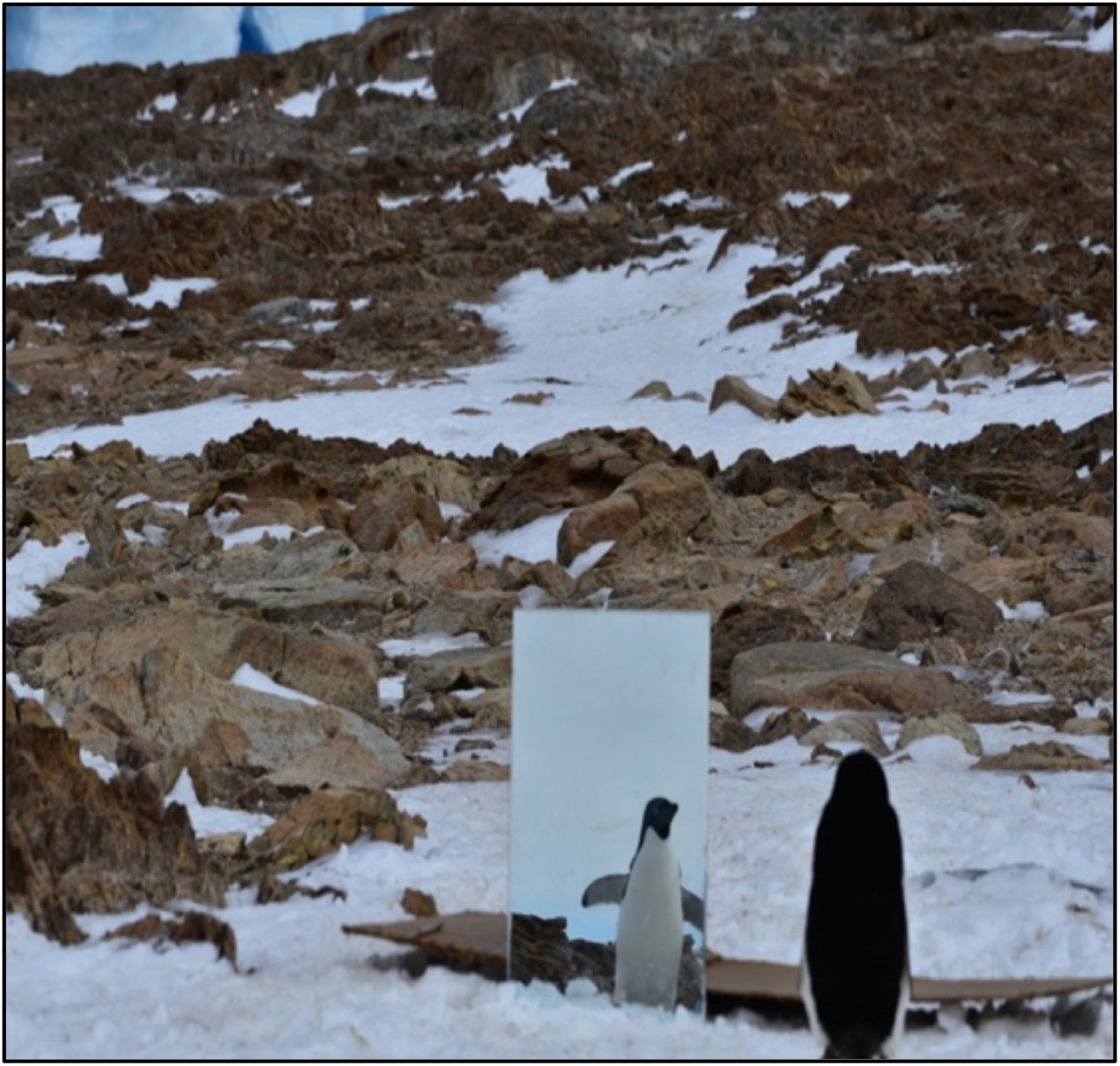
An Adélie penguin intently observing their own image during a group-behaviour test on Svenner Island, East Antarctica

#### 2.3.2 The modified mirror test, the hidden-head test and the coloured-bib test

Twelve, randomly selected, adult Adélie penguins, living in the rookery on Svenner Island, were employed in this set of experiments, which involved three experimental paradigms (Table 1). Presumably, the study individuals had neither undergone any kind of training nor had had any previous exposure to mirrors. In contrast to most previous studies on nonhuman species, we conducted these self-recognition tests in the natural habitat of the penguins, without subjecting them to any kind of captivity or prolonged, restrictive experience. We also believe that the moulting season, in which we conducted the experiments, may have provided extra alertness to the subject penguins.

**Table 1.** Details of the modified mirror test, hidden-head test and the coloured-bib test, conducted independently on twelve adult Adélie penguins on Svenner Island, East Antarctica

| Serial number | Date | Details of the experiment | Duration of the initial control phase | Duration of the later experimental phase |
| --- | --- | --- | --- | --- |
| Modified mirror test |  |  |  |  |
| 1. | 6 February 2020 | – | 7 sec | 42 sec |
| 2. | 6 February 2020 | – | 3 sec | 54 sec |
| 3. | 7 February 2020 | – | 5 sec | 1.14 min |
| Hidden-head test |  |  |  |  |
| 1. | 6 February 2020 | – | 6 sec | 10.54 min |
| 2. | 7 February 2020 | – | 44 sec | 3.57 min |
| 3. | 7 February 2020 | – | 5 sec | 2.18 min |
| 4. | 7 February 2020 | – | 5 sec | 4.18 min |
| Coloured-bib test |  |  |  |  |
| 1. | 6 February 2020 | Green bib | 6 sec | 6.31 min |
| 2. | 7 February 2020 | Red bib | 55 sec | 5.02 min |
| 3. | 7 February 2020 | Yellow bib | 6 sec | 0.40 min |
| 4. | 7 February 2020 | Yellow bib | 6 sec | 9.07 min |
| 5. | 7 February 2020 | Red bib | 6 sec | 8.15 min |

The first experimental paradigm, the modified mirror test, was conducted independently on three individual penguins in two three-sided, closed enclosures, constructed using cardboard sheets, with approximate dimensions of 68 × 100 cm and 68 × 40 cm respectively (Figure 3). The former, bigger enclosure was placed around an active adult penguin while the smaller enclosure was used to either to close the larger one or expand it by 40 cm, as and when required. Two glass mirrors – each 54 × 40 cm in dimension – were placed on the two opposite sides of the large enclosure to ensure that the three subject penguins confronted their mirror images within the enclosure. The penguins were released from their cardboard enclosures as soon as the experiments were concluded.

**Figure 3.**
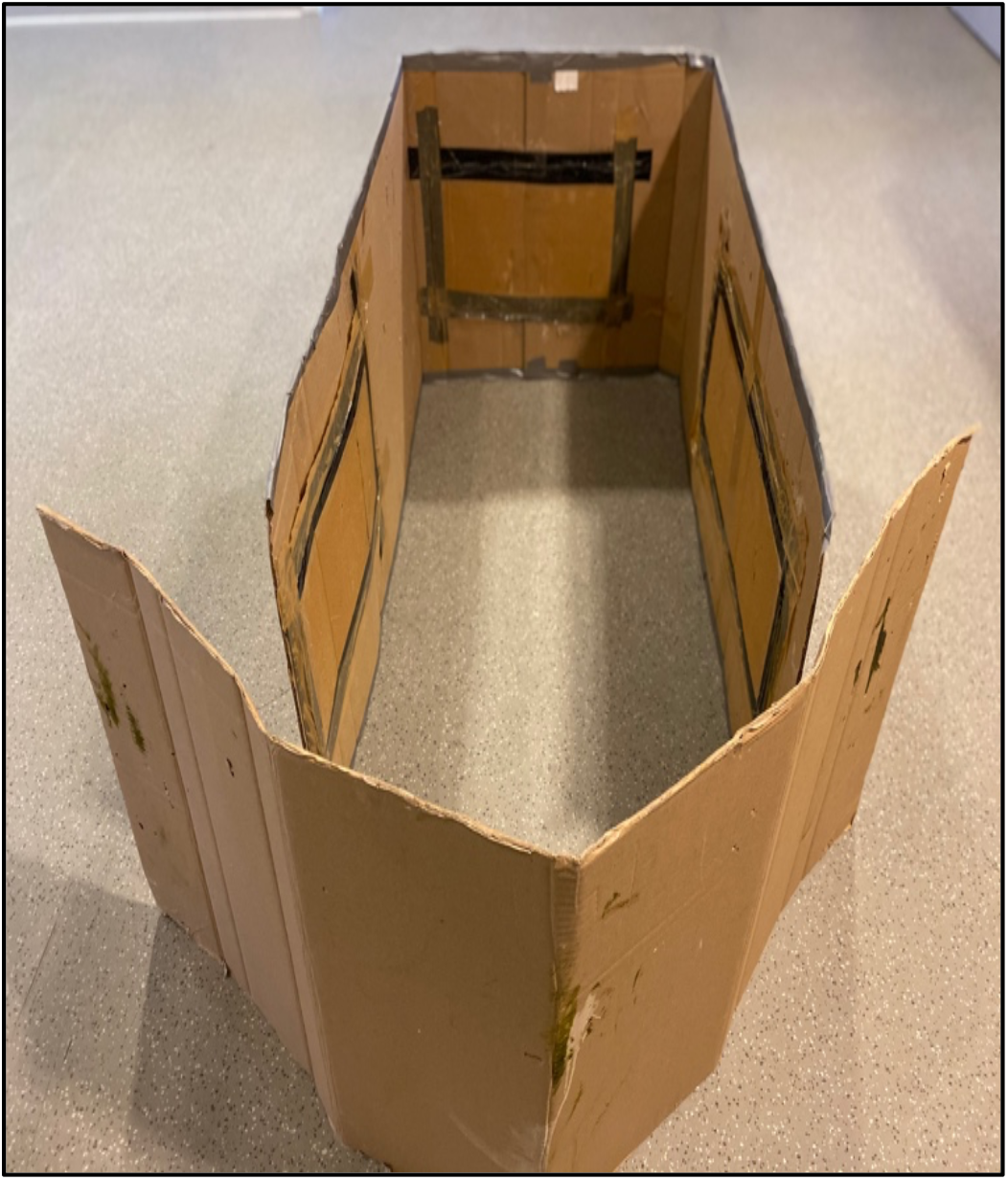
The cardboard enclosure that was used for all the three experimental paradigms in this study

Each experiment consisted of an initial control phase, usually lasting several seconds, when the penguins occupied the enclosure without any mirror (Table 1). The subsequent experimental phase consisted of the actual mirror test, when the two mirrors were placed in the enclosure and the subject penguins were observed to spend variable, but relatively longer, periods of time in front of the mirrors, apparently examining their images (Table 1). It should be noted that the control phase in each of these experiments was not extended for prolonged periods of time to ensure that the penguins did not become too restless, as this could potentially interfere with their subsequent performance in the later experimental phase of the trial.

The second experimental paradigm, the hidden-head test, which was conducted on four individual penguins, was virtually identical to the modified mirror test, but had a circular, green, paper disc, of diameter 14 cm, pasted on the front mirror in the larger enclosure, during the later experimental phase or the actual mirror test, to obstruct the reflection of the heads and upper body parts of the subject penguins. The disc was, however, absent during the initial control phase of each of these experiments although the mirrors were present. The behaviour of the subject penguins was observed in front of either mirror during the initial control phase and that in front of the mirror with the sticker during the later experimental phase, both phases lasting for different periods of time, as earlier (Table 1). Again, as before, all the subject penguins were released from the enclosures as soon as the tests were completed.

The final experimental paradigm, the coloured-bib test, was conducted independently on five adult penguins. Each of the subject individuals had a small bib of a certain colour, randomly chosen, fastened around their neck during the experimental phase and their behaviour in front of either of the two mirrors observed for different lengths of time, during the experimental phase of the test (Table 1). It should be noted that the way the bibs were attached did not obscure the contours of their bodies, as could be visualised in their mirror images. The penguins were also observed during the initial control phase of the experiments although they did not have their bibs attached during this phase of the test. Finally, the bibs were removed from the penguins as soon as the experiments were concluded and, as with the previous two sets of experiments, the subject penguins released without any further delay.

### 2.4 Research ethics clearance

All the procedures and protocols, followed in this study, was in strict accordance with ATCM Resolution 4 (2019) Annex of the Scientific Committee on Antarctic Research / SCAR’s Code of Conduct for the Use of Animals for Scientific Purposes in Antarctica, also referred to as “the Code of Conduct”.

## 3. Results and Discussion

### 3.1 The group-behaviour test

A waddle of twelve, healthy, adult Adélie penguins, living in the rookery on Svenner Island, was examined for their responses to the self-images that were generated in the experimental mirror, during one trial of the group-behaviour test. Several individuals from the waddle appeared to be simultaneously attracted to their images and stood relatively still, staring intently at the images for several seconds each, but making no attempts to either touch their images or reach out behind the mirror.

Similar behaviour was recorded in four individual adult penguins, belonging to different waddles, which were also observed to gaze at their images, while remaining relatively motionless, during four independent trials of the group-behaviour test (Figure 2). On all these occasions, the penguins—either as a group or individually—spent a range of 11 to 16 min in front of their respective images. Again, they did not direct any tactile behaviour towards their images in any of these experiments.

### 3.2 The modified mirror test, the hidden-head test and the coloured-bib test

In the three modified mirror tests conducted, the subject penguins displayed some restlessness and paced rapidly within their respective enclosures during the initial control phase, when no mirror was present in the enclosure. This agitation, however, significantly abated when the mirrors were introduced into the enclosure, and the penguins were exposed to their mirror images. Their focus immediately shifted to an exploration of their self-images (Figure 4). The penguins made rapid movements of their heads, flippers or of their bodies, some of which appeared to be gestures. Many of these movements and gestures were rapidly repeated but strikingly, the visual attention of all the subject penguins was firmly on their images during the entire timespan of their performance.

**Figure 4.**
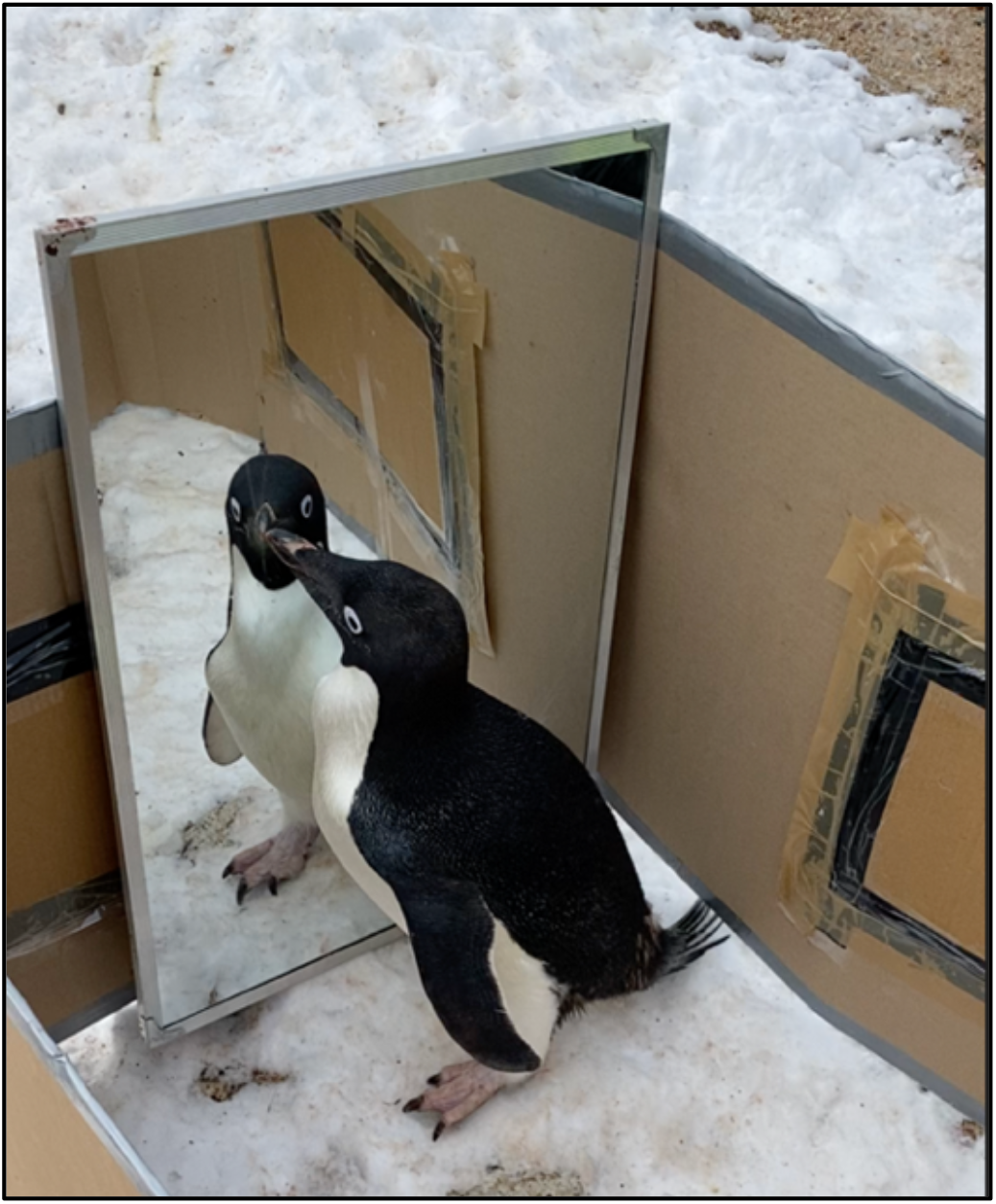
An Adélie penguin gazing intently at their image during a modified mirror test

Our observation that the penguins stared intently at the simultaneous body movements of their mirror images, as they proceeded to move their heads or other body parts, coupled with a singular lack of any kind of tactile reaching out or aggression directed towards the mirror images, suggested to us that the individuals possibly did not consider the images to represent other conspecific individuals. Whether the subject penguins recognised the images in the mirror to represent themselves, however, continues to remain an open question. Following Gallup Jr.’s original argument, such deliberate body movements could, nevertheless, be facilitated by the penguins’ ability to process “proprioceptive information and kinesthetic feedback onto the reflected visual image so as to coordinate the appropriate visually guided movements *(of their own bodies)* via the mirror” (Gallup Jr. 1970, p 87, phrase in italics ours). Moreover, the behavioural sequences, displayed variably by the subject penguins, appeared to be unique within the scope of our observations, as we never saw them performed by any penguin on any other occasion, either solitarily or when communicating with conspecific individuals within their waddles in their natural environment.

The behavioural performance of the four subject penguins during the initial control phase of the hidden-head test, when they were able to see the image of their complete bodies in the mirror, was similar to that of the penguins in the experimental phase of the modified mirror test, described above. There was, however, a stark difference in their behaviour in the later experimental phase of the test, when they were unable to see the image of their heads and upper parts of their body; it now primarily consisted of an active pecking of the obstructive stickers, accompanied by frenetic body movements (Figure 5).

**Figure 5.**
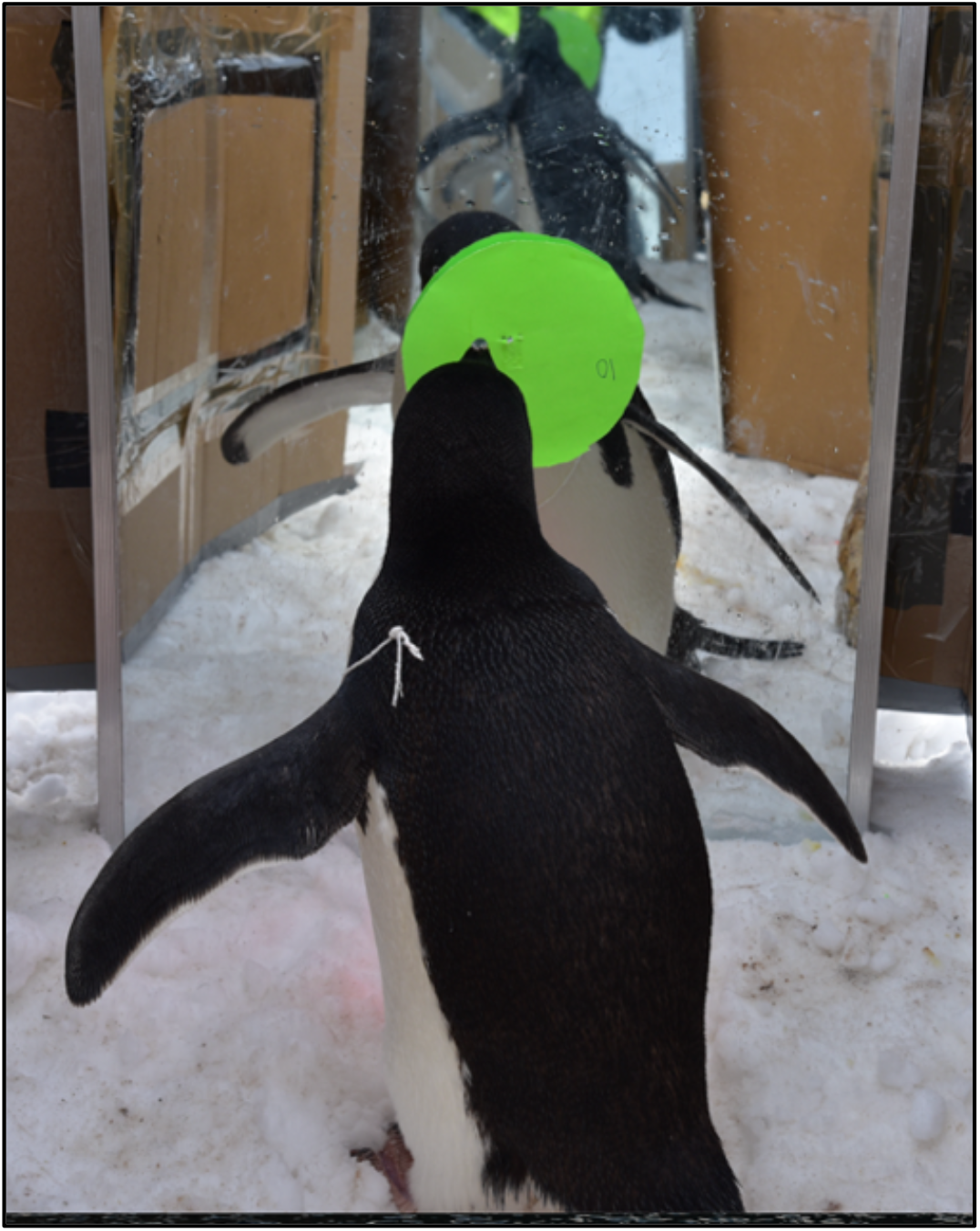
An Adélie penguin pecking on the circular disc, attached to a mirror, during a hidden-head test

We interpret the directed pecking of the subject penguins on the circular sticker as attempts to remove the obstruction, perhaps driven by an urge to restore the images that they had just seen earlier in the mirror. Furthermore, we speculate that such a behavioural motivation could indicate a restlessness that was expressed when they were unable to later see their faces in the mirror – a potential reflection of their underlying awareness of the self. We, however, admit that there could be alternative explanations, such as a discomfort generated by the failure to see the eyes of the mirror image (see, for example, Emery 2000 for the importance of the social gaze in nonhumans) for this behavioural response as well.

In the third set of field experiments that we carried out—the coloured-bib test—the penguins did not appear to exhibit any noticeable change in their apparently mirror-guided, rapid body movements, when the bibs were attached to their necks, from that displayed during the initial control phase of the experiment, without the bibs (Figure 6).

**Figure 6.**
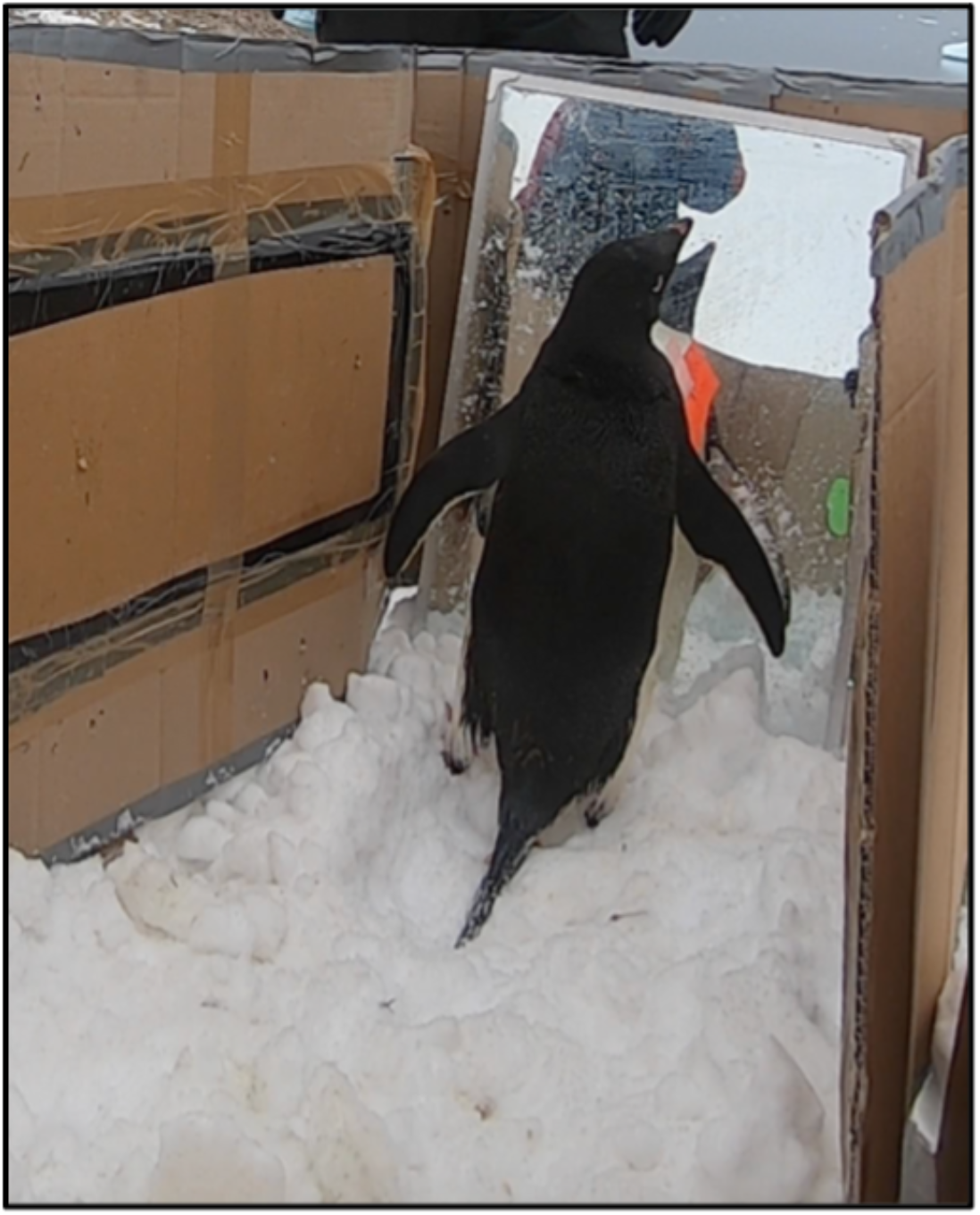
An Adélie penguin, with a yellow bib on the neck, inspecting their mirror image, during a coloured-bib test

These preliminary results remain equivocal about whether the penguins did perceive the coloured bibs around their necks during the tests and yet failed to respond to them or whether they failed to notice or see the bibs at all in their restless condition during the experiment. Anecdotal evidence suggests that certain penguin species may not see red although they appear to be sensitive to the violet, blue and green regions of the spectrum. Notwithstanding our first attempts to investigate whether our subject penguins would indeed explore extraneous visual stimuli on their bodies, when seen in their mirror images, such experiments should be better designed in the future.

We believe that being a social animal that lives in large mobile communities in the extreme environments of Antarctica and yet displays remarkable individuality in their social behaviour and communication (Penney 1968; Spurr 1974, 1975; Jouventin and Aubin 2002; Marks *et al*. 2010), natural selection may have putatively led to the awareness of their own bodies – and by extension, of their own selves – in Adélie penguins, as evidenced by the preliminary results that we have obtained from our simple mirror tests of self-awareness. Everyday interactions, at the individual level, with the co-inhabitants of one’s colony, we speculate, may have led to the appearance of recognition of their own bodies in this species – with the formation of distinct mental visual images of the different parts of their bodies, as has been seen earlier in primates (Dittrich 1994). Moreover, the remarkable capabilities of Adélie penguins to discriminate between conspecific individuals, especially within their large colonies (Jouventin and Aubin 2002; Marks *et al*. 2010), is another indication that the capacity to have a ‘self-concept’ of themselves as individuals may have been an important step during their socio-cognitive evolution (Griffin and Speck 2004). In a parallel set of studies on the Adélie penguin brain, we propose to investigate synaptic densities and other neuronal features in certain brain areas, which may reveal the possible evolution of such socio-cognitive capacities in these remarkably communal birds (Kandel 2001; Dastidar *et al*. 2022).

Finally, we hypothesise that Adélie penguins, given their intrinsic ability to immerse themselves in socially complex, networked lives within communal rookeries, may possess a sense of self-identity and subjective self-awareness, which characterises most, if not all, complex social species and which culminates in the most sophisticated form of symbolic self-awareness, apparently the hallmark of the human species alone (Sinha 2017). The ability of different nonhuman species to react variously to their mirror images may thus depend on their individual social dispositions and lifestyles. In primate groups, for example, a “qualitative psychological difference” has been noticed, following prolonged exposure to mirrors, and attributable to early social experiences (Gallup Jr. 1977, p 87). Although not investigated to any detail yet, we speculate that it is entirely possible that similar phenomena may exist in penguin species, including Adélie penguins, with their complex social lives within communal rookeries. Future studies, integrating the socioecology and cognitive ethology of penguins, may enable the testing of our hypothesis that complex social living, in which individual penguins coordinate certain aspects of their cooperative behaviour with conspecific individuals, while maintaining their independent decision-making capacities throughout their communal lives, could have led to the evolution of self-awareness, one of its manifestations being the ability to successfully negotiate the mirror self-recognition test that we tested them on (but see Brandl 2018).

## Acknowledgements

This project was financially and logistically supported by the National Centre for Polar and Ocean Research, Goa, India and the Ministry of Earth Sciences, Government of India, New Delhi. The authors express their sincere gratitude to M Ravichandran, Director, National Centre for Polar and Ocean Research, Goa, India for conceptual advancement of this work, and M Javed Beg, Vikas Gandhi, Punarbasu Choudhury and Michelle Fernandes for their help in designing the artefacts, used in the field tests, and in conducting the experiments. The authors would like to particularly acknowledge Captain Julius, a most capable helicopter pilot, for making the landings possible on the rough and rocky terrain of Svenner Island for the researchers to be able to successfully conduct their natural experiments.

## References

Bekoff M and Sherman PW. 2004. Reflections on animal selves. Trends in Ecology and Evolution, 19, 176–180

Brandl JL. 2018. The puzzle of mirror self-recognition. Phenomenology and the Cognitive Sciences, 17, 279–304

Brecht KF and Nieder A. 2020. Parting self from others: Individual and self-recognition in birds. Neuroscience and Biobehavioral Reviews, 116, 99–108

Brecht KF, Müller J and Nieder A. 2020. Carrion crows (Corvus corone corone) fail the mirror mark test yet again. Journal of Comparative Psychology, 134, 372–378

Buniyaadi A, Taufique SKT and Kumar V. 2020. Self-recognition in corvids: evidence from the mirror-mark test in Indian house crows (Corvus splendens). Journal of Ornithology, 161, 341–350

Cazzolla Gatti R, Velichevskaya A, Gottesman B and Davis K. 2021. Grey wolf may show signs of self-awareness with the sniff test of self-recognition. Ethology Ecology and Evolution, 33, 444–467

Croxall JP and Prince PA. 1979. Antarctic seabird and seal monitoring studies. Polar Record, 19, 573–595

Dastidar PG, Roy TS, Iyengar S, Kumaran S, Jacob T, Rawat A, Sivaperuman C, Ahluwalia BS and Sinha A. 2022. Stepwise construction of an anatomical atlas of the Adélie penguin brain: a multidisciplinary approach. Poster, SCAR 2022: Tenth SCAR Open Science Conference, online, August 2022. https://www.virtual.scar2022.org/eposter-detailes.php?token=OTM3

Dittrich W. 1994. How monkeys see others: discrimination and recognition of monkeys’ shape. Behavioural Processes, 33, 139–154

Emery NJ. 2000. The eyes have it: the neuroethology, function and evolution of social gaze. Neuroscience and Biobehavioral Reviews, 24, 581–604

Epstein R, Lanza RP and Skinner BF. 1981. “Self-awareness” in the pigeon. Science, 212, 695–696

Gallup Jr GG. 1970. Chimpanzees: self-recognition. Science, 167, 86–87

Gallup Jr GG. 1977. Self recognition in primates: a comparative approach to the bidirectional properties of consciousness. American Psychologist, 32, 329–338

Griffin DR and Speck GB. 2004. New evidence of animal consciousness. Animal Cognition, 7, 5–18

Jouventin P and Aubin T. 2002. Acoustic systems are adapted to breeding ecologies: individual recognition in nesting penguins. Animal Behaviour, 64, 747–757

Kandel E. 2001. The molecular biology of memory storage: a dialogue between genes and synapses. Science, 294, 1030–1038

Kohda M, Sogawa S, Jordan AL, Kubo N, Awata S, Satoh S, Kobayashi T, Fujita A and Bshary R. 2022. Further evidence for the capacity of mirror self-recognition in cleaner fish and the significance of ecologically relevant marks. PLoS Biology, 20, e3001529

Marks EJ, Rodrigo AG and Brunton DH. 2010. Ecstatic display calls of the Adélie penguin honestly predict male condition and breeding success. Behaviour, 147, 165–184

Penney RL. 1968. Territorial and social behavior in the Adélie penguin. In: Austin Jr OL (ed) Antarctic Bird Studies (Washington D C: American Geophysical Union), pp 83–131

Prior H, Schwarz A and Güntürkün O. 2008. Mirror-induced behavior in the magpie (Pica pica): evidence of self-recognition. PLoS Biology, 6, e202

Reiss D and Morrison R. 2017. Reflecting on mirror-self-recognition: a comparative view. In: Call J, Burghardt GM, Pepperberg IM, Snowdon CT and Zentall T (eds) APA Handbook of Comparative Psychology: Perception, Learning, and Cognition, Vol 2 (Washington D C: American Psychological Association), pp 745–763

Safina C. 2015. Beyond Words: What Animals Think and Feel (Basingstoke, UK: Picador)

Sinha A. 2017. Scio ergo sum: knowledge of the self in a nonhuman primate. Journal of the Indian Institute of Science, 97, 567–582

Soler M, Pérez-Contreras T and Peralta-Sánchez JM. 2014. Mirror-mark tests performed on jackdaws reveal potential methodological problems in the use of stickers in avian mark-test studies. PLoS One, 9, e86193

Spurr EB. 1974. Individual differences in aggressiveness of Adélie penguins. Animal Behaviour, 22, 611–616

Spurr EB. 1975. Communication in the Adelie penguin. In: Stonehouse B (ed) The Biology of Penguins. (London: Macmillan), pp 449–501

Vanhooland L-C, Bugnyar T, Massen JJM. 2020. Crows (Corvus corone ssp.) check contingency in a mirror yet fail the mirror-mark test. Journal of Comparative Psychology, 134, 158–169

